# Ancient recombination events between human herpes simplex viruses

**DOI:** 10.1101/093641

**Authors:** Sonia Burrel, David Boutolleau, Diane Ryu, Henri Agut, Kevin Merkel, Fabian H. Leendertz, Sébastien Calvignac-Spencer

**Author notes:** Address correspondence to Sebastien Calvignac-Spencer,.

## Abstract

Herpes simplex viruses 1 and 2 (HSV-1 and HSV-2) are seen as close relatives but also unambiguously considered as evolutionary independent units. Here, we sequenced the genomes of 18 HSV-2 isolates characterized by divergent UL30 gene sequences to further elucidate the evolutionary history of this virus. Surprisingly, genome-wide recombination analyses showed that all HSV-2 genomes sequenced to date comprise HSV-1 fragments. Using phylogenomic analyses, we could also show that two main HSV-2 lineages exist. One lineage is mostly restricted to sub-Saharan Africa while the other has reached a global distribution. Interestingly, only the worldwide lineage is characterized by ancient recombination events with HSV-1. Our findings highlight the complexity of HSV-2 evolution, a virus of likely zoonotic origin which later recombined with its human-adapted relative. They also suggest that co-infections with HSV-1 and 2 may have genomic and potentially functional consequences and should therefore be monitored more closely.

---

Human herpes simplex virus 1 and 2 (HSV-1 and 2; species *Human alphaherpesvirus 1* and *2*, genus *Simplexvirus,* subfamily *Alphaherpesvirus,* family *Herpesviridae,* order *Herpesvirales)* are closely related pathogens that usually cause recurrent mucosal lesions but are also, more rarely, responsible of severe diseases like neonatal morbidity and meningo-encephalitis. HSV-1 is essentially transmitted by oral-oral contacts and is responsible for oro-facial herpes while HSV-2 is primarily sexually transmitted and causes genital herpes. However, HSV-1 can be transmitted through oral sex and consequently cause genital herpes. Infection with HSV-1 is extremely common and affects 67% of the worldwide population (Looker, Magaret, May, et al. 2015). Infection with HSV-2 is less frequent but still reaches a global prevalence of 11% (Looker, Magaret, Turner, et al. 2015). Important regional variations in prevalence exist for both viruses, with the highest prevalence observed in Africa (87% and 31% for HSV-1 and HSV-2, respectively; Looker, Magaret, May, et al. 2015; Looker, Magaret, Turner, et al. 2015).

HSV-1 and HSV-2 show low overall genomic variability. HSV-1, however, has accumulated more variation than HSV-2 (maximum overall divergence 1.1% vs. 0.4%; Szpara, et al. 2014; Kolb, et al. 2015; Newman, et al. 2015). In conjunction, most HSV-2 open reading frames (ORF) exhibit little (if any) variability. Yet we recently discovered a HSV-2 variant (HSV-2v) characterized by unusually divergent UL30 gene sequences (maximum divergence of 2.4%; Burrel, et al. 2013; Burrel, et al. 2015). While HSV-2 genomic diversity is only weakly, if at all, geographically structured (Kolb, et al. 2015; Newman, et al. 2015), this new variant was also exceptional in that it was mainly recovered from sub-Saharan African individuals.

To clarify the evolutionary history of HSV-2v, we generated nearly complete genomes from 18 isolates originating in distinct patients. In brief, we extracted DNA from viral stock supernatants, prepared dual-indexed sequencing libraries and performed a single double-round in-solution hybridization capture on the pooled libraries using a target enrichment kit comprising baits covering the whole HSV-1 and HSV-2 genomes. The capture product was sequenced on a MiSeq^®^ platform (Illumina). In average, and after quality filtering, mapping and deduplication, we determined the sequence of 75% of the genomes.

We generated a concatenated alignment of 73 coding sequences (106,037 positions) that comprised of all new HSV-2v sequences and sequences from all georeferenced simplexvirus genomes determined from hominine hosts, including the only sequence derived from a chimpanzee (chimpanzee alpha-1 herpesvirus, ChHV; Severini, et al. 2013). We first ran phylogenomic analyses in a maximum likelihood (ML) framework. These analyses identified two main lineages: a previously unrecognized African lineage only comprising of HSV-2v sequences of which most originate from sub-Saharan Africa; and a worldwide lineage that was also detected in sub-Saharan Africa and includes some HSV-2v sequences (Figure 1).

**Figure 1.**
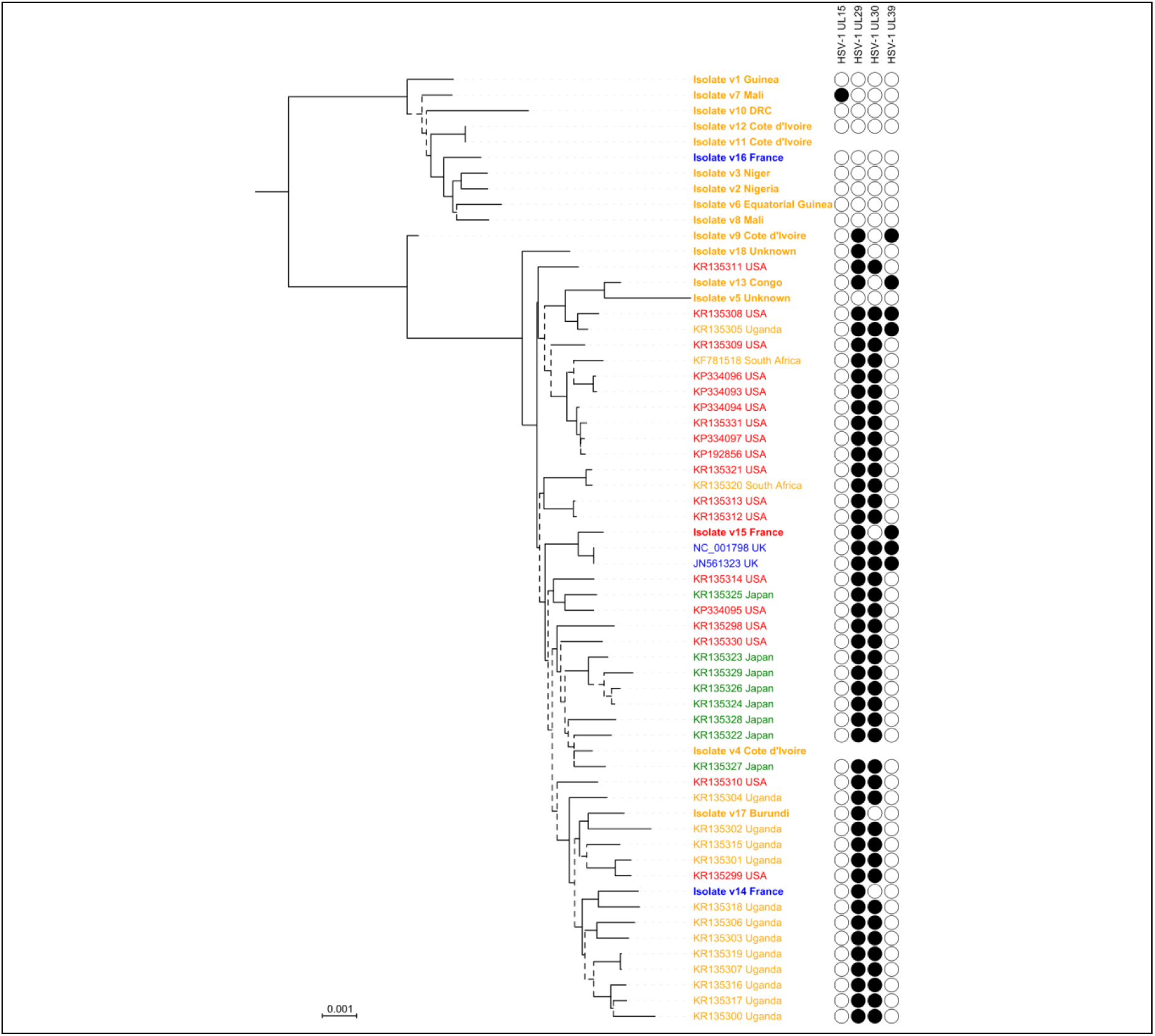
Phylogenomic analysis of HSV-2. This maximum likelihood tree was generated from an alignment of 106,037 positions comprising 61 sequences (including 43 publicly available genomes). Branch leaves are annotated with the accession number and country of origin of the virus; for sequences generated during this study isolate number and country name are in bold. The color code refers to the region of origin: orange for sub-Saharan Africa, blue for Europe, red for the Americas, green for Asia. Isolate v15 was obtained from a patient from Martinique (West Indies) and was therefore colored as originating in the Americas. Recombination profiles appear on the right side; filled circles indicate the presence of a HSV-1 fragment, empty circles their absence. Note that for 2 isolates (v4 and v11) we missed the data at the recombinant loci. Branch robustness was assessed using Shimodaira-Hasegawa-like approximate likelihood ratio tests (SH-like aLRT); branches supported by SH-like aLRT values < 0.95 are dashed. The scale is in substitutions per site. DRC: Democratic Republic of the Congo.

HSV-1 and HSV-2 are both prone to recombination although recombination events are seemingly more frequent in HSV-1 (Szpara, et al. 2014; Kolb, et al. 2015; Newman, et al. 2015). Therefore we also performed recombination analyses. This evidenced that members of the African and worldwide HSV-2 lineages recombined. For example, ML analyses performed on the five largest non-recombining fragments of the alignment showed that the HSV-2v isolate v9 (Côte d’Ivoire) belongs to the worldwide lineage for the first three fragments and to the African lineage for the last two (data not shown).

Unexpectedly, recombination analyses also identified interspecific recombination events between HSV-1 and HSV-2. We identified a minimum of 4 HSV-1 fragments recombined within 4 HSV-2 ORF: in UL15 (155 bp; positions 33,952-34,106 in the reference genome NC_001798), UL29 (397 bp; positions 60,003-60,399), UL30 (458 bp; positions 66,129-66,586) and UL39 (499 bp; positions 89,393-89,891). Of note, a close examination of the alignment may suggest 2 successive recombination events in UL29, with the HSV-2v isolate v13 (Congo) being an intermediate. Similarly, there may be some evidence of 2 partial reversal recombination events in UL39, where “native” HSV-2 may have recombined with HSV-2 already harboring a HSV-1 recombined fragment; the mark of these events can be seen in the sequences of the HSV-2v isolate v13 (Congo) and the HSV-2v isolate v15 (France) and two strains from the UK (accession numbers NC_001798 and JN561323), respectively.

To investigate the timing of the interspecific recombination events, we ran Bayesian analyses under a strict molecular clock model calibrated with the estimated divergence date of ChHV (Wertheim, et al. 2014) and annotated the trees with the recombination status at these 4 loci (Figure 1). This showed that the recombination events in UL29 and UL30 likely affected the ancestor of the worldwide lineage between 41 and 103 thousand years (ky) ago (95% highest posterior density - 95% HPD: 35-118 ky). The recombination events in UL39 and UL15 probably occurred later, respectively within the worldwide lineage less than 41 ky ago (95% HPD: 35-48 ky) and within the African lineage less than 20 ky ago (95% HPD: 16-25 ky).

Our results show that, contrary to common belief, mixed HSV-1/HSV-2 infections have led to natural gene flow from HSV-1 into HSV-2 genomes (Thiry, et al. 2005). Interestingly, we did not detect any gene flow in the opposite direction. Co-infection with HSV-1 and HSV-2 can occur in the anogenital zone on the condition that HSV-1 infection occurs prior to HSV-2 infection; HSV-2 infection seems to protect from further infection of the genitals with HSV-1 (Langenberg, et al. 1999; Looker and Garnett 2005). Such co-infections theoretically provide the opportunity of bidirectional gene flow. However, since HSV-1 causes fewer and milder recurrences in the anogenital zone than HSV-2, the transmission of recombinant HSV-1 comprising HSV-2 genome fragments can be expected to be very rare. This alone may explain the apparent unidirectional gene flow from HSV-1 into HSV-2.

Our findings also allow us to speculate on the evolution of HSV-2. The topology of the HSV-2 phylogeny is compatible with an African origin of HSV-2. In addition, the time to the most recent common ancestor of the worldwide lineage (excluding the recombinant HSV-2v isolate v9) is 41 ky (95% HPD: 35-48 ky), which shortly follows the single dispersal event from which all non-African human populations descended (Posth, et al. 2016). Therefore, it seems plausible that, following a putative transmission from chimpanzees to the human lineage (Wertheim, et al. 2014), HSV-2 diversified in African populations before a single lineage characterized by recombinant HSV-1 fragments (the worldwide lineage) accompanied the migration of *Homo sapiens* out of Africa. African HSV-2 belonging to the worldwide lineage may be the result of later contacts to non-African populations. We note here that this model, and more particularly the notion of an African origin of HSV-2, is also in line with the comparatively high HSV-2 prevalence observed today on the continent (Looker, Magaret, Turner, et al. 2015).

These results may also suggest a link between the recombination and evolutionary histories of HSV-2 since the recombinant fragments detected in UL29 and UL30 are hallmarks of the worldwide lineage. It is tempting to speculate that the success of this lineage was partly due to the respective early recombination events. Comparing the biological characteristics of strains from the worldwide and African lineages and/or engineered recombinants thereof may reveal their functional correlates (e.g. in terms of replication and transmission efficiency).

Finally, our results pinpoint that HSV-1/HSV-2 coinfections may have some public health relevance (since they already contributed to the genetic diversity of HSV-2 by allowing recombination events with HSV-1). Genomic monitoring will likely be the tool of choice to determine whether interspecific recombination is still an ongoing process and has any clinical implications.

